# The association between metabolite concentrations and wellbeing in adults

**DOI:** 10.1101/2024.07.12.603238

**Authors:** Natalia Azcona-Granada, Anne J.M.R. Geijsen, René Pool, Dirk H. M. Pelt, Meike Bartels

**Author notes:** Mailing address: Department of Biological Psychology, Vrije Universiteit Amsterdam, Amsterdam, Van der Boechorststraat 7, 1081 BT Amsterdam, the Netherlands, Corresponding author.

## Abstract

The biological complexity of wellbeing is studied from various perspectives, including genetics and epigenetics. However, there is a knowledge gap concerning other layers, such as metabolomics, which is dynamic and changes throughout life. This study explores the association between metabolites and wellbeing in a sample (N = 4748) drawn from the Netherlands Twin Register. A latent factor score for wellbeing was constructed based on: Quality of Life, Life Satisfaction, and Subjective Happiness. A total of 231 blood metabolites were analyzed using ^1^HNMR technique. Linear regression models were performed for each metabolite, while correcting for family clustering, relevant covariates, and multiple testing. None of the metabolites were significantly associated with wellbeing after multiple testing correction. Despite the lack of significant findings, the 34 metabolites with the lowest *p-*value (0.25) pointed to the same metabolic pathway: endogenous lipid metabolism. This pathway has previously been linked to wellbeing in a GWAS and associated with related phenotypes in other metabolomic studies. In conclusion, this study confirms the biological complexity of wellbeing and speculates on a potential role of lipids. Further research is needed to confirm these hypotheses.

## Introduction

Wellbeing (WB) is defined by the World Health Organization as a positive state experienced by individuals and societies, determined by social, economic, and environmental factors ^1^. In light of major demographic trends (e.g. aging populations, increasing longevity, decreasing birth rates, increasing inequality, and unforeseen pandemics), building and maintaining WB is one of the most important societal challenges. People who feel well, function better, are less susceptible to mental illness, and thus are better able to retain competitive advantage and expand human potential ^2–8^. Different conceptualizations of wellbeing exist, the most common distinction is between subjective wellbeing (SWB) and psychological wellbeing (PWB), also known as hedonic and eudaimonic wellbeing, respectively ^9–11^. The current study focuses, due to data availability, on SWB, defined as the cognitive and affective evaluation of one’s life.

WB is a complex phenotype influenced by various factors, including effects at different molecular biological levels ^12^. Biological complexity can be simplified in layers, each of them with different functions. Generally, these layers are denoted *omics*, which we can define as, amongst others, genomics, epigenomics, transcriptomics, proteomics, and metabolomics. They involve, respectively, the DNA/genes, methylation marks, transcripts, proteins, and metabolites. So each omics layer comprises different molecule types that allow for the specific functioning of it.

Some of them are very stable across life, such as the genome, since in healthy conditions, the genome remains the same regardless of for example time or exposure, cell type, or cell division cycle^13^. The role of genetics in WB has been previously studied with promising results ^14,15^. For example, a Genome-Wide Association Study found 304 significant independent SNPs associated with WB ^14^. The polygenic score for WB assessed in another study by Baselmans et at. (2019) explained between 0.62 and 0.76 of the variance observed in WB ^15^.

Other omics layers are subjective to changes by the environment or other conditions, e.g. epigenomics, transcriptomics, or metabolomics. The role of epigenomics in WB has also been studied. An Epigenome-Wide Association Study for WB reported also two significant sites after Bonferonni correction ^16^. Given these differences in characteristics, each omics layer can provide unique information about a trait of interest. For example, the metabolome that summarizes a given metabolic state of the body ^17^. Metabolites are molecules involved in or produced by biological processes, including digestion and other biochemical reactions ^17^ and the fact they are dynamic offers an interesting target for biological studies, intervention, and facilitating the implementation of more personalized medicine approaches ^18,19^.

Knowledge about the influence of omics layers beyond genomics or epigenomics on WB is currently lacking. A potential positive association between the neurotransmitter serotonin and WB (i.e., hedonic well-being) was revealed in a systematic review, but evidence on the role of other small molecules – including metabolites – was inconclusive ^20^. In addition, the precise interactions among these metabolites within the context of WB are not well understood and require further investigation.

Despite the absence of metabolomics studies for WB, interesting results have been observed in related phenotypes such as perceived stress and mental health. Noerman et al. showed associations between phosphatidylcholines and several parameters indicating stress, and found changes in a group labelled as “unknown class of lipids” over time to be correlated with physiological and psychological markers of stress ^21^. Jia et al. studied individuals with high and low depression and assessed cognitive skills and serum lipid profile (total cholesterol, triglycerides, HDL cholesterol, and LDL cholesterol) ^22^. In the low depression group, they found that the lower the LDL levels, the higher the cognitive scores. This may indicate that LDL could negatively affect cognitive skills, at least for individuals with low depressive symptoms. In addition, Bot et al.^23^ also reveals a distinct profile of circulating lipid metabolites linked to depression in a large-scale meta-analysis.

Summarizing, the aforementioned studies of Noerman et al.^21^, Jia et al.^22^ and Bot et al.^23^ combined with the meta-analysis results from de Vries et al.^20^ suggest that studying the association between metabolites in the context of wellbeing is of interest to better understand the interplay between metabolite concentrations and differences in WB. Therefore, we aim to test whether metabolites are associated with WB to increase our understanding of their biological interpretation concerning WB.

## Material and methods

This study was preregistered at the Open science framework platform (https://osf.io/4s2mu/).

### Participants

Participants included in this study are part of the Netherlands Twin Register (NTR), a population-based national register of twins and their families with extensive data collection on, amongst other things, mental health, personality, lifestyle, and demographics ^24^. Participants have been recruited since 1986, after which questionnaires have been collected every 2-3 years. In addition to questionnaires, biological samples were collected between 2004 and 2010. The questionnaires, biological sampling, processing, and storage are described in previously published work ^24,25^. In this study, we utilized surveys 6 (2006-2007), 8 (2009-2010), 10 (2013-2014), and 14 (2019-2020) to select a wellbeing measure optimally linked to the biological sample collection.

The data used in this study were obtained from the NTR after meeting the requirements posited by the NTR Data Management and the NTR Data Sharing Committee and obtaining their permission. All procedures performed in studies involving human participants were in accordance with the ethical standards of the institutional and/or national research committee and with the 1964 Helsinki Declaration. Data collection was approved by the Central Ethics Committee on Research Involving Human Subjects of the University Medical Centers Amsterdam. Informed consent was obtained from all individual participants included in the study. The ethical approval numbers are survey 6: 2001-069 (17-10-2001); 8: 2008-244 (01-12-2008); 10: 2011-334 (12-10-2011), 2012-433 (26-02-2013); 14: 2018-389 (25-07-2018) and for the biobank: 2003-180.

A sub-sample from the NTR biobank (N = 4748) was selected with available metabolomics and wellbeing data. The mean survey age was 40.6 [standard deviation (*SD*) 14.0, range 14-82 years], for 18 participants age was missing. The mean biological data age was 40.6 [standard deviation (*SD*) 13.6, range 18-79 years]. The ratio of men/women was 1536 (32%) /3210 (68%); 2 individuals did not report their sex.

### Measures

#### Wellbeing

Wellbeing was assessed as a continuous trait with three measures, i.e. Quality of Life with the Cantril Ladder (QoL; ^26^), the Satisfaction With Life Scale (SWLS; ^27^), and the Subjective Happiness Scale (SHS; ^28^). Quality of Life was assessed with the Cantril ladder, which requires participants to indicate the step of the ladder at which they place their lives in general on a 10-point scale (10 being the best possible life, and 1 the worst possible life) ^26^. The SWLS consists of five items answered on a seven-point scale ranging from 1=“strongly disagree” to 7=“strongly agree” ^27^. An example item is: “I am satisfied with my life”. The SHS consists of four items which rated on a seven-point scale ranging from 1=“strongly disagree” to 7=“strongly agree” ^28^. An example item is “On the whole I am a happy person”.

The different wellbeing measures were used in various surveys. QoL was part of survey 8, 10 and 14, SWLS was part of all surveys, while SHS was part of surveys 6, 8, 14. For SWLS and SHS the items – 5 items for SWLS, 4 for SHS – were summed to arrive at a sum score for each measure. Where necessary, inversion of item scores was done to ensure that for each variable a higher score indicated higher wellbeing levels. The sum scores for QoL, SWLS and SHS were first transformed into Z-scores to avoid scaling issues when creating the latent factor score.

In each wave, a single factor score for wellbeing was built on all available wellbeing measures (differences in number of available measurements per wave did not affect factor scores, see Supplementary information) using structural equation modeling ^29^. We used full information maximum likelihood when constructing the factor score, which can handle missing values on indicator variables. However, to limit the number of missing values per participant, only those with at least two out of three measurements per wave were included. To reduce bias, a transformation was used so that the covariance matrix of the estimated factor scores matched the model-based latent factor covariance matrix ^30,31^.

#### Metabolomics data

231 metabolic markers were quantified from plasma samples using high-throughput proton nuclear magnetic resonance spectroscopy (^1^H-NMR) metabolomics (Nightingale Health Ltd, Helsinki, Finland; formerly Brainshake Ltd.). This method provides simultaneous quantification of lipids, lipoprotein subclass profiling with lipid concentrations within 14 subclasses, fatty acid compositions, and various low molecular weight metabolites including amino acids, ketone bodies, and glycolysis-related metabolites in molar concentration units ^32^. Variables representing percentages of metabolites instead of the actual concentrations were removed to avoid collinearity. The final dataset contained 148 metabolites and 9 ratios, shown in Supplementary Table 1. All the variables were scaled to allow comparisons of findings. Four metabolites – FALen, UnsatDeg, UnSat, Gln – were missing in most of the participants (31, 31, 88, and 100%, respectively) resulting in a final set of 144 metabolites and 9 ratios for analyses.

The Nightingale Health 1-HNMR platform protocol is as follows: Before the NMR measurements, samples are mixed with a sodium phosphate buffer (75 mmol/L Na_2_HPO_4_ in 80%/20% H_2_O/D_2_O, pH 7.4; including also 0.08% sodium 3-(trimethylsilyl)propionate-2,2,3,3-d4 and 0.04% sodium azide) and moved to the NMR tubes. A PerkinElmer JANUS Automated Workstation with an 8-tip dispense arm with Varispan is responsible for the liquid handling. Samples are transferred to 96-well plates, with every plate includes two quality control (QC) samples (consistency of quantifications), a serum mimic and a mixture of two low-molecular–weight metabolites (performance automated liquid handler and spectrometer). The laboratory setup combines a Bruker AVANCE III 500 MHz and Bruker AVANCE III HD 600 MHz spectrometers, both with the SampleJet robotic sample changer. The 500 MHz spectrometer is furnished with a selective inverse room temperature probe head, the 600 MHz spectrometer is furnished with a cryogenically cooled triple resonance probe head (CryoProbe Prodigy TCI). The 500 MHz and 600 MHz spectrometers can both automatically collect the lipoprotein (LIPO) and low-molecular-weight metabolites (LMWM) with standardized parameters. After spectroscopy, the samples are processed manually with a standardized lipid extraction procedure with multiple extraction steps using an Integra Biosciences VIAFLO 96 channel electronic pipette. These lipid extracts are again analyzed, in full automation, with the 600 MHz spectrometer with a standard parameter set. Initial data processing is handled by the computers controlling the spectrometers. Processing includes automated phasing and Fourier transformation of the NMR spectra. In house-algorithms located on a centralized server process the spectral data further, which includes baseline removal, background control, checking for missing or extra peaks, and spectral area-specific signal alignments. Regression modelling is performed to quantify the molecular data for those spectral areas that pass QC.

#### Data selection

To reduce bias of changes over time, we aimed to keep the time between the biological sample collection and the survey as short as possible. Therefore, the wellbeing latent score with the least time between phenotype assessment and biological sample collection was selected. The final sample consists of 4,748 individuals, with 2,198 individuals with data from survey 6, 2,318 with data from survey 8, 199 from survey 10 and 33 from survey 14. The average time difference between the survey data and the biological data collection (time difference) was 2.49 years (SD = 0.89).

### Covariates

All models were adjusted for BMI, smoking status, fasting status, medication use (lipid-lowering, statin, hypertension, beta blocker, cardiac or antidepressive medication), time difference, and batch effects since these are variables that could affect the metabolite levels and/or measurements^33^.

The mean BMI of the sample was 24.7 (SD = 4.1). Smoking status was reported from the participants as no smokers (2515; 53%), former smokers (1307; 28%), and current smokers (912; 19%), while 14 individuals did not report smoking information. In total, 4479 (96%) samples were taken in a fasting state (data were missing for 72 participants), and 422 (9%) were using medication at the moment of the biological data collection.

To test whether the confounders included in the model of interest influence the variability in metabolite concentrations (collectively), Principal Component Partial R-square was used ^34^. Since we cannot create a separate model for each metabolite with specific confounders, this approach allowed us to determine the overall relationship between the confounders and the metabolite concentrations.

### Statistical Analyses

The removal of the variables for the metabolite percentages as well as the merging of the metabolite measures with the covariates were performed in Python 3 and the statistical analyses were performed in R, version 4.3.2. The main packages used were *pandas* for Python and *foreign* ^35^*, lavaan* ^36^*, gee* ^37^*, pcpr2* ^38^ for the statistical analyses in R.

Multiple linear regression in generalized estimating equation (GEE) models were used to determine the associations between the wellbeing score as the independent variable and concentrations of metabolites as dependent variables. GEE models allow to correct for non-independence of data introduced by twin status within families ^39^. All models were adjusted for BMI, smoking status, fasting status, medication use, time difference, and batch effect.

After correction for multiple testing, using false discovery rate (FDR) according to the Benjamin-Hochberg procedure ^40,41^, a *p*-value (pFDR)□<□0.05 was considered statistically significant.

## Results

The GEE models showed 52 significant associations between metabolites concentrations and wellbeing before FDR correction (*p* < 0.05). After the correction, no significant association remained (*pFDR* < 0.05). The *p*-values (before and after correction) as well as the direction of the effect – determined by the standardized β – are shown in Figure 1.

**Figure 1.**
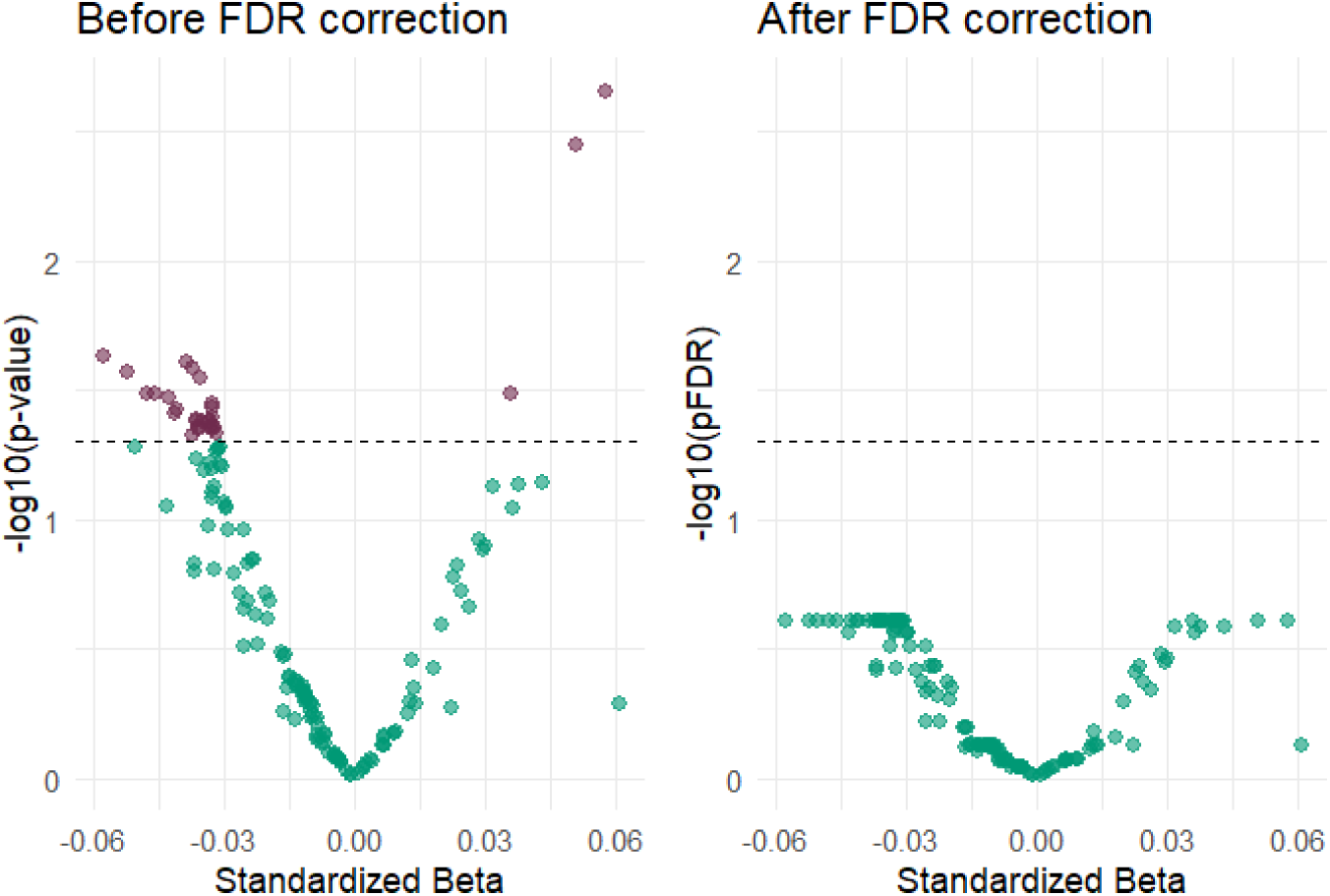
P-value of the metabolites before and after FDR correction. Associations between metabolite concentrations and wellbeing before and after multiple testing correction. P value vs standardized beta before (left) and after (right) FDR correction. -log10 of P value in the Y axis and standardized beta on the X axis. A threshold is set at 0.05 with a dotted line. Significant p values are shown in purple.

Given the absence of significant metabolites, we focused on the lowest *p*-value metabolites associated with wellbeing. After the p-value correction, 34 metabolites presented the same p-value (*pFDR* = 0.246), which was also the closest p-value to the significance threshold (0.05). The biological classification for all metabolites, and results of the GEE models for the 34 metabolites with top pFDR are described in Supplementary Table 1 and Table 2, respectively. Out of these 34 variables, 28 are of the class of lipoproteins, 2 are cholesterols, 2 glycerides and phospholipids, 1 is a lipoprotein, and 1 is of class “fluid balance”. Standardized beta values and their CI are represented in Figure 2.

**Figure 2.**
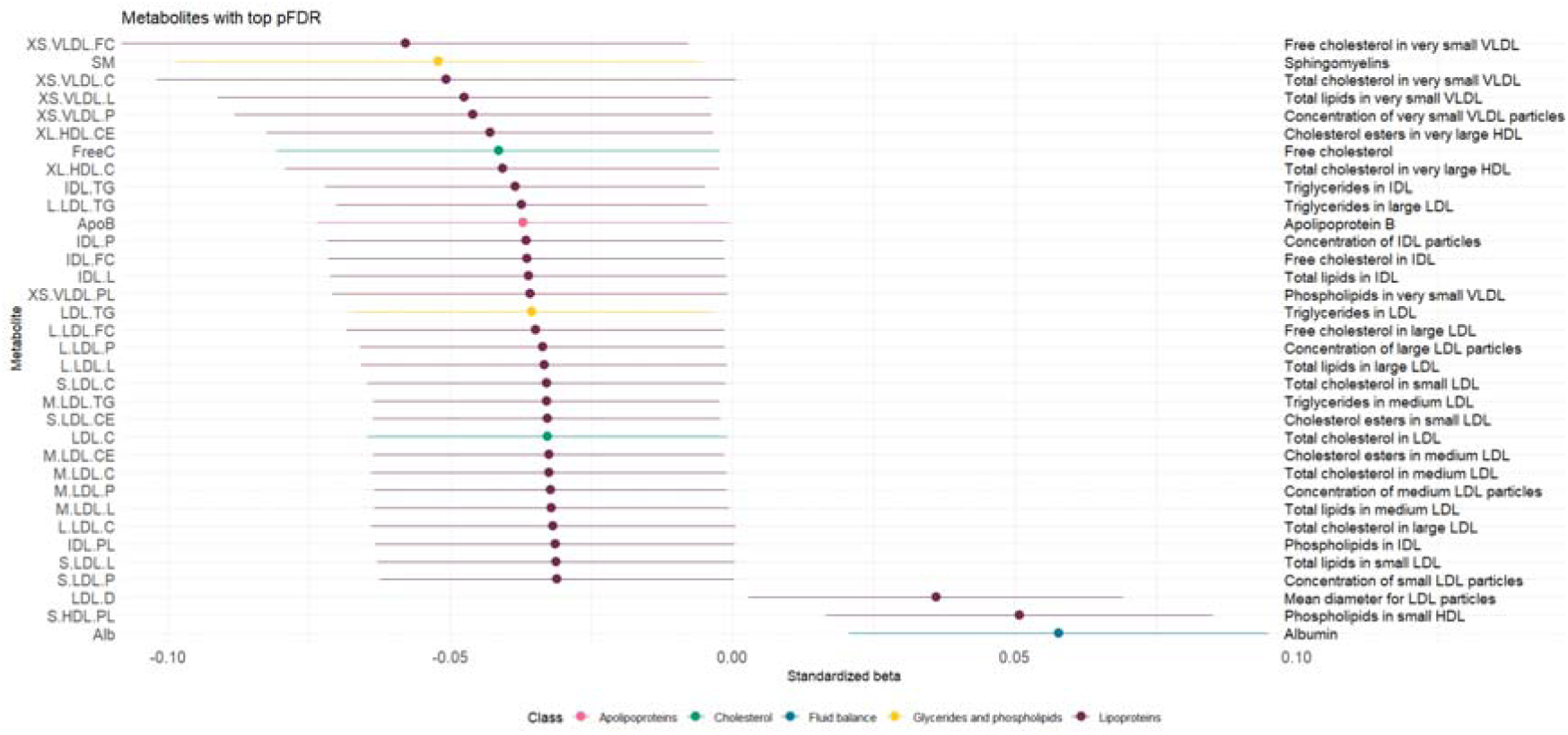
Forest plot of the metabolites with the lowest p-value after correction. Forest plot of the metabolites with the lowest p-value after correction (0.2456) from higher beta to lower colored by metabolite class (legend on the bottom). On the right the description of the metabolite. Note: when correction of p-value, only the p-value is corrected. For this reason, the metabolites are not significant even though their confidence intervals do not cross the 0 value.

The PCA R^2^ analysis showed that 21.35% of the variability in metabolite concentrations – all taken together – was explained by the confounders in the model of interest; BMI, smoking status, fasting status, medication use, time difference between biological and survey collection, and batch effect. The variability explained by each of the covariates as well as the total, is shown in Figure 3.

**Figure 3.**
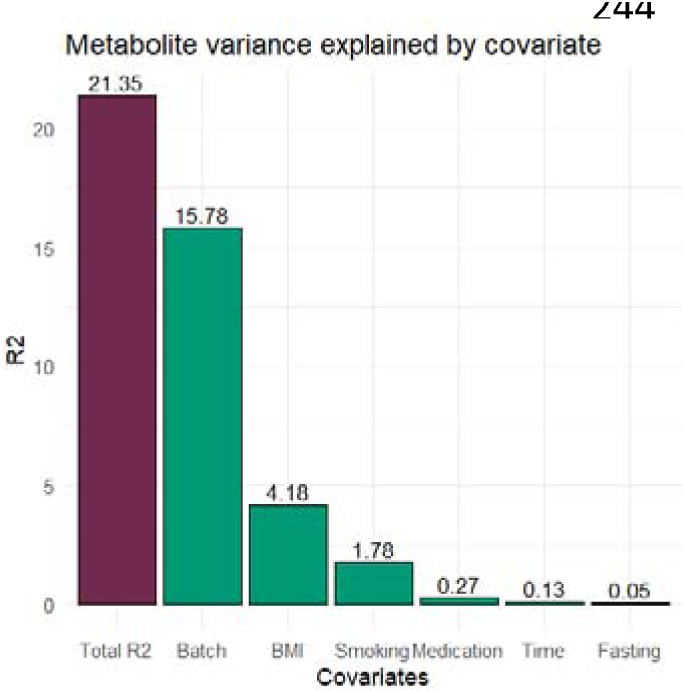
Metabolite variance explained by covariate. Metabolite variance explained by the confounders in the model of interest from higher to lowest. In purple the total R^2^ explained by the confounders and in green the covariate-specific R^2^.

## Discussion

We studied the relationship between metabolites and wellbeing in a sample drawn from the Netherlands Twin Register (NTR). Our results showed no significant associations between metabolite concentrations or ratios and wellbeing after multiple testing corrections, but provided an interesting proof of concept of the complexity of the biology of wellbeing.

As far as we know, our study is the first focusing on the association between metabolites and wellbeing, which prevents us from directly comparing our results with previous work. In this study, from the original 148 metabolites, 34 metabolites were reported with the same lowest *p*-value after FDR correction (*pFDR* = 0.24563). These are mainly (apo)lipoproteins and cholesterol molecules. Given the fact that they were not statistically significant, we hypothesize a broad explanation of their role in the biology of wellbeing, regardless of the size of the molecules and the specific parameters for each molecule.

All the 34 metabolites mentioned above take part in the endogenous lipid metabolism: the lipoproteins (Very Low Density Lipoprotein (VLDL), Intermediate Density Lipoprotein (IDL), Low Density Lipoprotein (LDL), and High Density Lipoprotein (HDL)), the apolipoproteins (Apo B) and phospholipids (Triglycerides and Sphingomyelins) closely related to them, the free cholesterol (Free Cholesterol), and even albumin (in charge of lipid transportation in blood ^42^) can be connected to this pathway, illustrated in Figure 4. In general, lipids are essential for cell structures and to generate energy in the cell ^17^.

**Figure 4.**
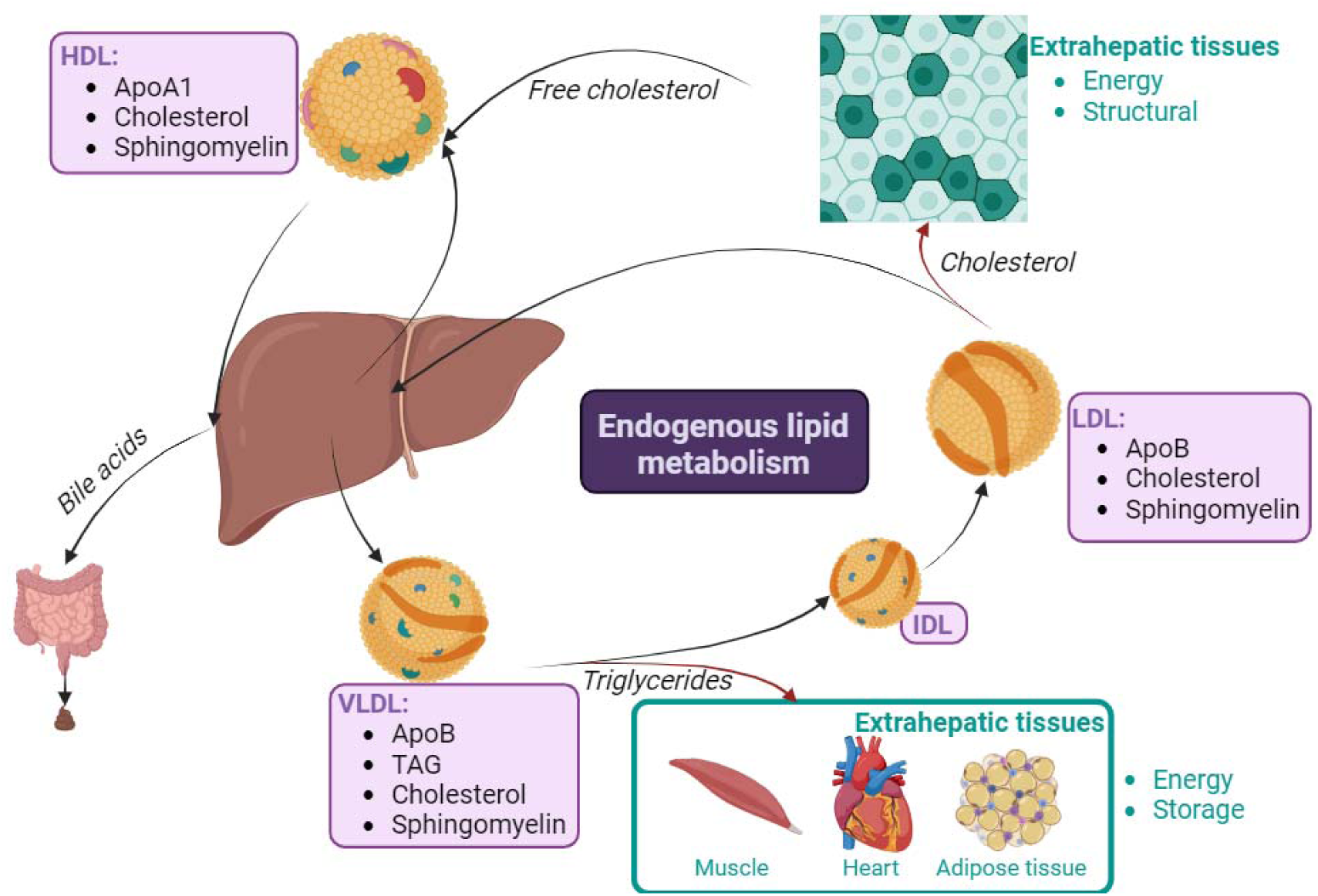
Endogenous lipid metabolism scheme. The liver synthesizes Triglycerides and cholesterol, which circulate in VLDL lipoproteins containing ApoB and sphingomyelin. These VLDLs transport lipids to tissues, where they release Triglycerides and transform into IDLs and LDLs, carrying cholesterol. LDLs return cholesterol to the liver or tissues, but excess LDLs can oxidize, leading to atheroma formation, earning them the label “bad cholesterol.” Conversely, HDLs, containing ApoA1 and sphingomyelin, gather excess cholesterol and transport it back to the liver for disposal, earning them the moniker “good cholesterol”.

An hypothesis previously stated in the literature is that a healthier lifestyle is associated with a happier life ^43^. “Bad” cholesterol – transported by LDL – and the associated molecules can have a negative effect on the cardiovascular system and are associated with a poorer lifestyle ^44^. While the “good” cholesterol molecules – transported by HDL and associated with a healthy and balanced lifestyle – can have a positive effect on the cardiovascular system ^44^. In line with this hypothesis, we found the βs for LDL to be negative while the βs for HDL were positive, which implies that the lower the LDL or the higher the HDL parameters, the higher wellbeing levels are reported. However, as the results were not significant they are suggestive, warranting caution in their interpretation and additional research for replication.

This potential effect of lipid metabolism on wellbeing is in line with previous results in other wellbeing omics, such as genomics. Baselmans et al. found in a Genome-Wide Association Study (GWAS) for wellbeing a significant effect of a SNP (*rs10838629*) associated with the LDL receptor (LDL receptor-related protein 4 (*LRP4*)) ^14^. In another study from Turley et al. (2018), there are also some significant SNPs for WB that are associated with lipid-related molecules, such as LDL receptor-related protein 1B (*LRP1B*, *rs10197004*) and Apolipoprotein A1 (*APOA1, rs61905145*) ^45^. Therefore, the pathway standing out in our results is in line with the same pathway reported in previous genetic studies for wellbeing.

Metabolomics results of other phenotypes closely related to wellbeing are worth discussing here. Noerman et al. found associations between phosphatidylcholines and several parameters indicating subjective stress. They also observed that changes in an unknown class of lipids over time correlate with physiological and psychological markers of stress ^21^. Unfortunately, we did not include phosphatidylcholines or so-called “unknown class of lipids” parameters and, therefore, results cannot be compared directly.

As noted, Jia et al., suggested that serum lipid profile (total cholesterol, triglycerides, HDL cholesterol, and LDL cholesterol) may be linked directly to self-rated depression and cognitive performance ^22^. For the low depression group, they found that the lower the LDL levels, the higher the cognitive scores. Although the phenotype is not exactly similar to the current study hypothesizes, an effect of the cholesterol pathway is observed. LDL, specifically, is present in the top metabolites subset observed in the present study and with the same (potential) direction of effect. The pattern observed in the discussed studies investigating metabolites in relation to wellbeing and mental health, is that lipids may play a role in wellbeing or related phenotypes compared to the other large molecule groups – proteins, nucleic acids, and carbohydrates. Thus, future research should focus on lipids in studies with a bigger sample size to unravel the potential relationship between lipid concentrations and wellbeing.

There are notable limitations in this study that require consideration. Firstly, the biological samples were not collected on the same day as the surveys were filled out with a time difference between 2 and 13 years (average of 2.49 years, SD = 0.89) with the biological sampling, potentially introducing bias in identifying metabolites associated with wellbeing. Unlike the genome, metabolite levels are highly dynamic. Even though wellbeing is assumed to be relatively stable across time ^46–51^, differences between the time of biological sample and survey collection may have influenced our results, since metabolite concentrations change over time and conditions ^18,19^. Sensitivity analyses with a time difference below 5 years, resulting in a reduction in sample size of 4.93% – 4514 individuals remained –, did not result in notable differences with the main analyses (see Supplementary Analyses and Supplementary Table 3). In addition, the PCA *R*^2^ showed that only 0.13% of the variance in the total set of metabolites was explained by the time difference. Still, we cannot rule out that our results are partly influenced by differences in data collection between individuals. Secondly, this is a cross-sectional study – i.e., unable to assess causality – only able to reveal associations. Third, the focus on lipids in the available data may overlook important insights from other metabolite classes that were not covered with the used technique. Lastly, surveys 6 and 10 had only two measurements available instead of 3, which meant that the factor scores of survey 6 and 10 could have been less reliable than the others. However, we found that the correlations between all the factor scores were not significantly different from each other (Supplementary information), meaning that there was no bias introduced by the survey used.

Conversely, this study presents several strengths. The sample size is substantial for a study of this nature (N = 4,748). The variables included in the models as covariates were also found to be important to include, given the results of the analysis. The PCA-R2 analysis showed that 21% of the variability observed in the metabolites is accounted for by the covariates. This underlines the importance of correcting for them, as done in this study.

A challenge with testing for associations between metabolites and an outcome is that many metabolites are correlated. Having several, correlated measurements for the same molecule or set of molecules – in combination with multiple testing correction – might have acted to our disadvantage. Alternatively, by reducing the number of subtypes per molecule, the multiple-testing burden could be lowered, although it is hard to decide a priori which measurements should be taken. Rather than selecting fewer measurements per molecule, data reduction techniques could be applied to find subgroups of metabolites associated with the outcome of interest ^52^. However, data reduction techniques always imply losing information and may complicate the biological interpretability of obtained results ^53^.

Future investigations should focus on lipids, examining lipoproteins regardless of their subtype or other parameters should be considered, preferably in an even larger sample. Alternatively, one could include other classes of metabolites, not covered by the used technique in this study, to confirm that the recommended focus on lipids is appropriate. Additionally, data from other biological layers should be included to do an integrative multi-omics framework in the future, promising a deeper understanding of their interactions in determining wellbeing.

In conclusion, in this study, it is speculated that lipids may play a role in the metabolism of wellbeing, but more research including other omics is necessary to disentangle the complex biology of wellbeing.

## Acknowledgments

We warmly thank all participating twins in the Netherlands Twin Register who dedicated part of their time to make research possible as well as everyone involved in the collection of the data and data management. This study is funded by an NWO-VICI grant (VI.C.211.054, PI Bartels). Prof Bartels is furthermore funded by an ERC Consolidator Grant (WELL-BEING; grant 771057). Data collection for this study has been funded by NWO large investment grant (NTR: 480-15-001/674), ZonMW Addiction program (31160008), ERC Consolidator Grant (WELL-BEING; grant 771057), NWO-MW 904-61-193, NWO 575-25-006, Genotype/phenotype database for behavior genetic and genetic epidemiological studies (ZonMw Middelgroot 911-09-032); the Biobank-based integrative omics study (BIOS) funded by BBMRI-NL (NWO projects 184.021.007 and 184.033.111).

## Ethical statement

All procedures performed in studies involving human participants were in accordance with the ethical standards of the institutional and/or national research committee and with the 1964 Helsinki declaration. Data collection was approved by the Central Ethics Committee on Research Involving Human Subjects of the University Medical Centers Amsterdam. Informed consent was obtained from all individual participants included in the study. Ethical approval numbers are as follows: survey 6: 2001-069 (17-10-2001); 8: 2008-244 (01-12-2008); 10: 2011-334 (12-10-2011), 2012-433 (26-02-2013); 14: 2018-389 (25-07-2018) and for the biobank: 2003-180.

## Declarations

The authors report there are no competing interests to declare

### Code availability

https://github.com/NataliaAG99/MetabolomicsWellbeing_NTR

### Data availability

Being part of a national prospective cohort study, Netherlands Twin Register data cannot be made publicly available for privacy reasons, but they are available for legitimate researchers via the data access procedure (https://ntr-data-request.psy.vu.nl/). The research data collected by the Netherlands Twin Registry (NTR) are pseudonymized, annotated, and stored in the NTR Repository. This is a secure database that is only accessible to our data managers. Metadata (i.e. variable names, labels, counts etc.) can be consulted by researchers in the NTR Data Showcase (https://ntr-data-request.psy.vu.nl/data-showcase.html). The Data Showcase allows researchers to create a list of variables and export it for use in a Data Sharing Request (https://ntr-data-request.psy.vu.nl/submitting-a-data-sharing-request.html). Researchers with an approved data sharing request who abide by the rules of the European General Data Protection Regulation will receive temporary access to the NTR data for their own research projects. More information on our data-sharing procedures and associated costs can be found https://ntr-data-request.psy.vu.nl/data-sharing-procedures.html. The datasets used and/or analyzed during the current study are available from the corresponding author upon reasonable request.

