## Supplementary Material for "The association between metabolite concentrations and wellbeing in adults"

Supplementary Information. Wellbeing score in the surveys.

To test the robustness of the factor score with respect to missing assessments of wellbeing in two of the four surveys (survey 6 and 10 only had two assessments while survey 8 and 14 had an alternative score, also based on two measures) was calculated for survey 8 and 14 and correlated with the score built on three measurements. This robustness check returned a correlation between the 2-measures score and the 3-measures score of 0.94 for survey 8 and 0.97 for survey 14. Thus, using two or three measurements to construct a factor score did not influence the factor scores in a significant way supporting the robustness of the method to construct the score.

|  | *Survey 6* | *Survey 8* | *Survey 10* | *Survey 14* |
| --- | --- | --- | --- | --- |
| *Survey 6* | 1 | 0.55 | 0.49 | 0.54 |
| *Survey 8* |  | 1 | 0.62 | 0.55 |
| *Survey 10* |  |  | 1 | 0.62 |
| *Survey 14* |  |  |  | 1 |

***Supplementary Table.*** *Correlations between the factor scores for wellbeing between surveys.*

Supplementary Table 1. Classification of the metabolites included in the analysis.

| Name | Description | Main class |
| --- | --- | --- |
| XXL-VLDL-P | Concentration of chylomicrons and extremely large VLDL particles | Lipoprotein |
| XXL-VLDL-L | Total lipids in chylomicrons and extremely large VLDL | Lipoprotein |
| XXL-VLDL-PL | Phospholipids in chylomicrons and extremely large VLDL | Lipoprotein |
| XXL-VLDL-C | Total cholesterol in chylomicrons and extremely large VLDL | Lipoprotein |
| XXL-VLDL-CE | Cholesterol esters in chylomicrons and extremely large VLDL | Lipoprotein |
| XXL-VLDL-FC | Free cholesterol in chylomicrons and extremely large VLDL | Lipoprotein |
| XXL-VLDL-TG | Triglycerides in chylomicrons and extremely large VLDL | Lipoprotein |
| XL-VLDL-P | Concentration of very large VLDL particles | Lipoprotein |
| XL-VLDL-L | Total lipids in very large VLDL | Lipoprotein |
| XL-VLDL-PL | Phospholipids in very large VLDL | Lipoprotein |
| XL-VLDL-C | Total cholesterol in very large VLDL | Lipoprotein |
| XL-VLDL-CE | Cholesterol esters in very large VLDL | Lipoprotein |
| XL-VLDL-FC | Free cholesterol in very large VLDL | Lipoprotein |
| XL-VLDL-TG | Triglycerides in very large VLDL | Lipoprotein |
| L-VLDL-P | Concentration of large VLDL particles | Lipoprotein |
| L-VLDL-L | Total lipids in large VLDL | Lipoprotein |
| L-VLDL-PL | Phospholipids in large VLDL | Lipoprotein |
| L-VLDL-C | Total cholesterol in large VLDL | Lipoprotein |
| L-VLDL-CE | Cholesterol esters in large VLDL | Lipoprotein |
| L-VLDL-FC | Free cholesterol in large VLDL | Lipoprotein |
| L-VLDL-TG | Triglycerides in large VLDL | Lipoprotein |
| M-VLDL-P | Concentration of medium VLDL particles | Lipoprotein |
| M-VLDL-L | Total lipids in medium VLDL | Lipoprotein |
| M-VLDL-PL | Phospholipids in medium VLDL | Lipoprotein |
| M-VLDL-C | Total cholesterol in medium VLDL | Lipoprotein |
| M-VLDL-CE | Cholesterol esters in medium VLDL | Lipoprotein |
| M-VLDL-FC | Free cholesterol in medium VLDL | Lipoprotein |
| M-VLDL-TG | Triglycerides in medium VLDL | Lipoprotein |
| S-VLDL-P | Concentration of small VLDL particles | Lipoprotein |
| S-VLDL-L | Total lipids in small VLDL | Lipoprotein |
| S-VLDL-PL | Phospholipids in small VLDL | Lipoprotein |
| S-VLDL-C | Total cholesterol in small VLDL | Lipoprotein |
| S-VLDL-CE | Cholesterol esters in small VLDL | Lipoprotein |
| S-VLDL-FC | Free cholesterol in small VLDL | Lipoprotein |
| S-VLDL-TG | Triglycerides in small VLDL | Lipoprotein |
| XS-VLDL-P | Concentration of very small VLDL particles | Lipoprotein |
| XS-VLDL-L | Total lipids in very small VLDL | Lipoprotein |
| XS-VLDL-PL | Phospholipids in very small VLDL | Lipoprotein |
| XS-VLDL-C | Total cholesterol in very small VLDL | Lipoprotein |
| XS-VLDL-CE | Cholesterol esters in very small VLDL | Lipoprotein |
| XS-VLDL-FC | Free cholesterol in very small VLDL | Lipoprotein |
| XS-VLDL-TG | Triglycerides in very small VLDL | Lipoprotein |
| IDL-P | Concentration of IDL particles | Lipoprotein |
| IDL-L | Total lipids in IDL | Lipoprotein |
| IDL-PL | Phospholipids in IDL | Lipoprotein |
| IDL-C | Total cholesterol in IDL | Lipoprotein |
| IDL-CE | Cholesterol esters in IDL | Lipoprotein |
| IDL-FC | Free cholesterol in IDL | Lipoprotein |
| IDL-TG | Triglycerides in IDL | Lipoprotein |
| L-LDL-P | Concentration of large LDL particles | Lipoprotein |
| L-LDL-L | Total lipids in large LDL | Lipoprotein |
| L-LDL-PL | Phospholipids in large LDL | Lipoprotein |
| L-LDL-C | Total cholesterol in large LDL | Lipoprotein |
| L-LDL-CE | Cholesterol esters in large LDL | Lipoprotein |
| L-LDL-FC | Free cholesterol in large LDL | Lipoprotein |
| L-LDL-TG | Triglycerides in large LDL | Lipoprotein |
| M-LDL-P | Concentration of medium LDL particles | Lipoprotein |
| M-LDL-L | Total lipids in medium LDL | Lipoprotein |
| M-LDL-PL | Phospholipids in medium LDL | Lipoprotein |
| M-LDL-C | Total cholesterol in medium LDL | Lipoprotein |
| M-LDL-CE | Cholesterol esters in medium LDL | Lipoprotein |
| M-LDL-FC | Free cholesterol in medium LDL | Lipoprotein |
| M-LDL-TG | Triglycerides in medium LDL | Lipoprotein |
| S-LDL-P | Concentration of small LDL particles | Lipoprotein |
| S-LDL-L | Total lipids in small LDL | Lipoprotein |
| S-LDL-PL | Phospholipids in small LDL | Lipoprotein |
| S-LDL-C | Total cholesterol in small LDL | Lipoprotein |
| S-LDL-CE | Cholesterol esters in small LDL | Lipoprotein |
| S-LDL-FC | Free cholesterol in small LDL | Lipoprotein |
| S-LDL-TG | Triglycerides in small LDL | Lipoprotein |
| XL-HDL-P | Concentration of very large HDL particles | Lipoprotein |
| XL-HDL-L | Total lipids in very large HDL | Lipoprotein |
| XL-HDL-PL | Phospholipids in very large HDL | Lipoprotein |
| XL-HDL-C | Total cholesterol in very large HDL | Lipoprotein |
| XL-HDL-CE | Cholesterol esters in very large HDL | Lipoprotein |
| XL-HDL-FC | Free cholesterol in very large HDL | Lipoprotein |
| XL-HDL-TG | Triglycerides in very large HDL | Lipoprotein |
| L-HDL-P | Concentration of large HDL particles | Lipoprotein |
| L-HDL-L | Total lipids in large HDL | Lipoprotein |
| L-HDL-PL | Phospholipids in large HDL | Lipoprotein |
| L-HDL-C | Total cholesterol in large HDL | Lipoprotein |
| L-HDL-CE | Cholesterol esters in large HDL | Lipoprotein |
| L-HDL-FC | Free cholesterol in large HDL | Lipoprotein |
| L-HDL-TG | Triglycerides in large HDL | Lipoprotein |
| M-HDL-P | Concentration of medium HDL particles | Lipoprotein |
| M-HDL-L | Total lipids in medium HDL | Lipoprotein |
| M-HDL-PL | Phospholipids in medium HDL | Lipoprotein |
| M-HDL-C | Total cholesterol in medium HDL | Lipoprotein |
| M-HDL-CE | Cholesterol esters in medium HDL | Lipoprotein |
| M-HDL-FC | Free cholesterol in medium HDL | Lipoprotein |
| M-HDL-TG | Triglycerides in medium HDL | Lipoprotein |
| S-HDL-P | Concentration of small HDL particles | Lipoprotein |
| S-HDL-L | Total lipids in small HDL | Lipoprotein |
| S-HDL-PL | Phospholipids in small HDL | Lipoprotein |
| S-HDL-C | Total cholesterol in small HDL | Lipoprotein |
| S-HDL-CE | Cholesterol esters in small HDL | Lipoprotein |
| S-HDL-FC | Free cholesterol in small HDL | Lipoprotein |
| S-HDL-TG | Triglycerides in small HDL | Lipoprotein |
| VLDL-D | Mean diameter for VLDL particles | Lipoprotein |
| LDL-D | Mean diameter for LDL particles | Lipoprotein |
| HDL-D | Mean diameter for HDL particles | Lipoprotein |
| Serum-C | Serum total cholesterol | Cholesterol |
| VLDL-C | Total cholesterol in VLDL | Cholesterol |
| Remnant-C | Remnant cholesterol (non-HDL, non-LDL -cholesterol) | Cholesterol |
| LDL-C | Total cholesterol in LDL | Cholesterol |
| HDL-C | Total cholesterol in HDL | Cholesterol |
| HDL2-C | Total cholesterol in HDL2 | Cholesterol |
| HDL3-C | Total cholesterol in HDL3 | Cholesterol |
| EstC | Esterified cholesterol | Cholesterol |
| FreeC | Free cholesterol | Cholesterol |
| Serum-TG | Serum total triglycerides | Glycerides and phospholipids |
| VLDL-TG | Triglycerides in VLDL | Glycerides and phospholipids |
| LDL-TG | Triglycerides in LDL | Glycerides and phospholipids |
| HDL-TG | Triglycerides in HDL | Glycerides and phospholipids |
| TotPG | Total phosphoglycerides | Glycerides and phospholipids |
| TG/PG | Ratio of triglycerides to phosphoglycerides | Glycerides and phospholipids |
| PC | Phosphatidylcholine and other cholines | Glycerides and phospholipids |
| SM | Sphingomyelins | Glycerides and phospholipids |
| TotCho | Total cholines | Glycerides and phospholipids |
| ApoA1 | Apolipoprotein A-I | Apolipoproteins |
| ApoB | Apolipoprotein B | Apolipoproteins |
| ApoB/ApoA1 | Ratio of apolipoprotein B to apolipoprotein A-I | Apolipoproteins |
| TotFA | Total fatty acids | Fatty acid |
| FALen | Estimated description of fatty acid chain length, not actual carbon number | Fatty acid |
| UnsatDeg | Estimated degree of unsaturation | Fatty acid |
| DHA | 22:6, docosahexaenoic acid | Fatty acid |
| LA | 18:2, linoleic acid | Fatty acid |
| FAw3 | Omega-3 fatty acids | Fatty acid |
| FAw6 | Omega-6 fatty acids | Fatty acid |
| PUFA | Polyunsaturated fatty acids | Fatty acid |
| MUFA | Monounsaturated fatty acids; 16:1, 18:1 | Fatty acid |
| SFA | Saturated fatty acids | Fatty acid |
| DHA/FA | Ratio of 22:6 docosahexaenoic acid to total fatty acids | Fatty acid |
| LA/FA | Ratio of 18:2 linoleic acid to total fatty acids | Fatty acid |
| FAw3/FA | Ratio of omega-3 fatty acids to total fatty acids | Fatty acid |
| FAw6/FA | Ratio of omega-6 fatty acids to total fatty acids | Fatty acid |
| PUFA/FA | Ratio of polyunsaturated fatty acids to total fatty acids | Fatty acid |
| MUFA/FA | Ratio of monounsaturated fatty acids to total fatty acids | Fatty acid |
| SFA/FA | Ratio of saturated fatty acids to total fatty acids | Fatty acid |
| Glc | Glucose | Glycolysis related |
| Lac | Lactate | Glycolysis related |
| Cit | Citrate | Glycolysis related |
| Ala | Alanine | Amino acid |
| Gln | Glutamine | Amino acid |
| His | Histidine | Amino acid |
| Ile | Isoleucine | Amino acid |
| Leu | Leucine | Amino acid |
| Val | Valine | Amino acid |
| Phe | Phenylalanine | Amino acid |
| Tyr | Tyrosine | Amino acid |
| Ace | Acetate | Ketone bodies |
| AcAce | Acetoacetate | Ketone bodies |
| bOHBut | 3-hydroxybutyrate | Ketone bodies |
| Crea | Creatinine | Fluid balance |
| Alb | Albumin | Fluid balance |
| Gp | Glycoprotein acetyls, mainly a1-acid glycoprotein | Inflammation |

***Supplementary Table 1.*** *Name of the metabolite (Name); description of the metabolite (Description); and main class (MainClass).*

Supplementary Table 2. Descriptives of the 34 metabolites with the lowest *p*-value after FDR correction.

| **Metabolite** | **Description** | **β_St_** | **se** |  | **CI** |  | | **P-value** | | **pFDR** |
| --- | --- | --- | --- | --- | --- | --- | --- | --- | --- | --- |
| XS-VLDL-P | Concentration of very small VLDL particles | -0.046 | 0.022 | -0.088 / -0.004 | | | 0.0329 | | 0.2456 | |
| XS-VLDL-L | Total lipids in very small VLDL | -0.047 | 0.022 | -0.091 / -0.004 | | | 0.0326 | | 0.2456 | |
| XS-VLDL-PL | Phospholipids in very small VLDL | -0.036 | 0.018 | -0.071 / -0.001 | | | 0.0450 | | 0.2456 | |
| XS-VLDL-C | Total cholesterol in very small VLDL | -0.051 | 0.026 | -0.102 / 0.001 | | | 0.0527 | | 0.2456 | |
| XS-VLDL-FC | Free cholesterol in very small VLDL | -0.058 | 0.026 | -0.108 / -0.008 | | | 0.0236 | | 0.2456 | |
| IDL-P | Concentration of IDL particles | -0.037 | 0.018 | -0.072 / -0.002 | | | 0.0409 | | 0.2456 | |
| IDL-L | Total lipids in IDL | -0.036 | 0.018 | -0.071 / -0.001 | | | 0.0444 | | 0.2456 | |
| IDL-PL | Phospholipids in IDL | -0.031 | 0.016 | -0.063 / 0.001 | | | 0.0539 | | 0.2456 | |
| IDL-FC | Free cholesterol in IDL | -0.036 | 0.018 | -0.071 / -0.001 | | | 0.0422 | | 0.2456 | |
| IDL-TG | Triglycerides in IDL | -0.038 | 0.017 | -0.072 / -0.005 | | | 0.0249 | | 0.2456 | |
| L-LDL-P | Concentration of large LDL particles | -0.034 | 0.016 | -0.066 / -0.001 | | | 0.0415 | | 0.2456 | |
| L-LDL-L | Total lipids in large LDL | -0.033 | 0.017 | -0.066 / -0.001 | | | 0.0440 | | 0.2456 | |
| L-LDL-C | Total cholesterol in large LDL | -0.032 | 0.016 | -0.064 / 0.001 | | | 0.0546 | | 0.2456 | |
| L-LDL-FC | Free cholesterol in large LDL | -0.035 | 0.017 | -0.068 / -0.001 | | | 0.0415 | | 0.2456 | |
| L-LDL-TG | Triglycerides in large LDL | -0.037 | 0.017 | -0.070 / -0.004 | | | 0.0260 | | 0.2456 | |
| M-LDL-P | Concentration of medium LDL particles | -0.032 | 0.016 | -0.063 / -0.001 | | | 0.0438 | | 0.2456 | |
| M-LDL-L | Total lipids in medium LDL | -0.032 | 0.016 | -0.063 / -0.001 | | | 0.0461 | | 0.2456 | |
| M-LDL-C | Total cholesterol in medium LDL | -0.032 | 0.016 | -0.064 / -0.001 | | | 0.0437 | | 0.2456 | |
| M-LDL-CE | Cholesterol esters in medium LDL | -0.033 | 0.016 | -0.064 / -0.001 | | | 0.0407 | | 0.2456 | |
| M-LDL-TG | Triglycerides in medium LDL | -0.033 | 0.016 | -0.064 / -0.002 | | | 0.0354 | | 0.2456 | |
| S-LDL-P | Concentration of small LDL particles | -0.031 | 0.016 | -0.062 / 0.000 | | | 0.0522 | | 0.2456 | |
| S-LDL-L | Total lipids in small LDL | -0.031 | 0.016 | -0.063 / 0.000 | | | 0.0531 | | 0.2456 | |
| S-LDL-C | Total cholesterol in small LDL | -0.033 | 0.016 | -0.065 / -0.001 | | | 0.0419 | | 0.2456 | |
| S-LDL-CE | Cholesterol esters in small LDL | -0.033 | 0.016 | -0.064 / -0.002 | | | 0.0370 | | 0.2456 | |
| XL-HDL-C | Total cholesterol in very large HDL | -0.041 | 0.020 | -0.079 / -0.002 | | | 0.0378 | | 0.2456 | |
| XL-HDL-CE | Cholesterol esters in very large HDL | -0.043 | 0.020 | -0.083 / -0.003 | | | 0.0337 | | 0.2456 | |
| S-HDL-PL | Phospholipids in small HDL | 0.051 | 0.017 | 0.017 / 0.085 | | | 0.0036 | | 0.2456 | |
| LDL-D | Mean diameter for LDL particles | 0.036 | 0.017 | 0.003 / 0.069 | | | 0.0328 | | 0.2456 | |
| LDL-C | Total cholesterol in LDL | -0.033 | 0.016 | -0.065 / -0.001 | | | 0.0450 | | 0.2456 | |
| FreeC | Free cholesterol | -0.041 | 0.020 | -0.081 / -0.002 | | | 0.0388 | | 0.2456 | |
| LDL-TG | Triglycerides in LDL | -0.035 | 0.016 | -0.067 / -0.004 | | | 0.0286 | | 0.2456 | |
| SM | Sphingomyelins | -0.052 | 0.023 | -0.098 / -0.006 | | | 0.0267 | | 0.2456 | |
| ApoB | Apolipoprotein B | -0.037 | 0.019 | -0.074 / 0.000 | | | 0.0474 | | 0.2456 | |
| Alb | Albumin | 0.058 | 0.019 | 0.021 / 0.095 | | | 0.0022 | | 0.2456 | |

***Supplementary Table 2.*** *Name of the metabolite (Metabolite); description of their classification (Description); standardized beta (β_St_); standard error (se); 95% confidence interval (CI); p-value (P-value) and p-value after FDR correction (pFDR).*

Supplementary Analyses. Regression analyses with a time difference of less than 5 years between the survey data and the biological data.

Excluding all participants for whom the time difference between the survey date and the biological sample collection date was more than five years we lost information for 234 individuals, retaining a sample of 4514 individuals. Of them, 2171 individuals had data from survey 6 while 2246 had data from survey 8 and only 97 individuals from survey 10 remained.

In this new analysis, 13 metabolites were significant before the FDR correction but no metabolites remained significant after the aforementioned FDR correction. When focusing on the metabolites with the lowest p-value (0.3036), the results showed similar standardized beta coefficients and *p*-values, except for Leucine and Valine, which were not present in the previous subgroup. These metabolites are shown in the table below.

| **Metabolite** | **Description** | **β_St_** | **se** | **CI** | **p-val** | **pFDR** |
| --- | --- | --- | --- | --- | --- | --- |
| XS-VLDL-P | Concentration of very small VLDL particles | -0.043 | 0.022 | -0.086 / 0.000 | 0.051 | 0.304 |
| XS-VLDL-L | Total lipids in very small VLDL | -0.044 | 0.023 | -0.088 / 0.001 | 0.053 | 0.304 |
| XS-VLDL-PL | Phospholipids in very small VLDL | -0.033 | 0.018 | -0.069 / 0.002 | 0.064 | 0.304 |
| XS-VLDL-FC | Free cholesterol in very small VLDL | -0.051 | 0.026 | -0.102 / -0.001 | 0.046 | 0.304 |
| IDL-P | Concentration of IDL particles | -0.035 | 0.018 | -0.070 / 0.000 | 0.052 | 0.304 |
| IDL-L | Total lipids in IDL | -0.034 | 0.018 | -0.070 / 0.001 | 0.057 | 0.304 |
| IDL-PL | Phospholipids in IDL | -0.030 | 0.016 | -0.062 / 0.002 | 0.068 | 0.304 |
| IDL-FC | Free cholesterol in IDL | -0.035 | 0.018 | -0.070 / 0.001 | 0.053 | 0.304 |
| IDL-TG | Triglycerides in IDL | -0.038 | 0.018 | -0.072 / -0.004 | 0.030 | 0.304 |
| L-LDL-P | Concentration of large LDL particles | -0.033 | 0.017 | -0.065 / 0.000 | 0.050 | 0.304 |
| L-LDL-L | Total lipids in large LDL | -0.032 | 0.017 | -0.065 / 0.000 | 0.053 | 0.304 |
| L-LDL-C | Total cholesterol in large LDL | -0.031 | 0.017 | -0.063 / 0.002 | 0.067 | 0.304 |
| L-LDL-FC | Free cholesterol in large LDL | -0.034 | 0.017 | -0.067 / 0.000 | 0.052 | 0.304 |
| L-LDL-TG | Triglycerides in large LDL | -0.037 | 0.017 | -0.071 / -0.004 | 0.029 | 0.304 |
| M-LDL-P | Concentration of medium LDL particles | -0.031 | 0.016 | -0.063 / 0.000 | 0.052 | 0.304 |
| M-LDL-L | Total lipids in medium LDL | -0.031 | 0.016 | -0.063 / 0.001 | 0.055 | 0.304 |
| M-LDL-C | Total cholesterol in medium LDL | -0.032 | 0.016 | -0.063 / 0.000 | 0.052 | 0.304 |
| M-LDL-CE | Cholesterol esters in medium LDL | -0.032 | 0.016 | -0.063 / 0.000 | 0.047 | 0.304 |
| M-LDL-TG | Triglycerides in medium LDL | -0.033 | 0.016 | -0.064 / -0.002 | 0.039 | 0.304 |
| S-LDL-P | Concentration of small LDL particles | -0.030 | 0.016 | -0.062 / 0.002 | 0.066 | 0.304 |
| S-LDL-L | Total lipids in small LDL | -0.030 | 0.016 | -0.062 / 0.002 | 0.067 | 0.304 |
| S-LDL-C | Total cholesterol in small LDL | -0.032 | 0.016 | -0.064 / 0.000 | 0.053 | 0.304 |
| S-LDL-CE | Cholesterol esters in small LDL | -0.032 | 0.016 | -0.063 / -0.001 | 0.045 | 0.304 |
| XL-HDL-C | Total cholesterol in very large HDL | -0.039 | 0.020 | -0.078 / 0.001 | 0.055 | 0.304 |
| XL-HDL-CE | Cholesterol esters in very large HDL | -0.040 | 0.021 | -0.081 / 0.001 | 0.054 | 0.304 |
| S-HDL-PL | Phospholipids in small HDL | 0.044 | 0.017 | 0.010 / 0.078 | 0.011 | 0.304 |
| LDL-D | Mean diameter for LDL particles | 0.036 | 0.017 | 0.003 / 0.070 | 0.035 | 0.304 |
| LDL-C | Total cholesterol in LDL | -0.032 | 0.016 | -0.064 / 0.001 | 0.054 | 0.304 |
| FreeC | Free cholesterol | -0.041 | 0.020 | -0.081 / -0.001 | 0.044 | 0.304 |
| LDL-TG | Triglycerides in LDL | -0.036 | 0.017 | -0.069 / -0.004 | 0.028 | 0.304 |
| SM | Sphingomyelins | -0.049 | 0.024 | -0.096 / -0.002 | 0.039 | 0.304 |
| ApoB | Apolipoprotein B | -0.036 | 0.019 | -0.073 / 0.002 | 0.060 | 0.304 |
| Leu | Leucine | 0.033 | 0.018 | -0.003 / 0.069 | 0.069 | 0.304 |
| Val | Valine | 0.040 | 0.018 | 0.005 / 0.076 | 0.027 | 0.304 |
| Alb | Albumin | 0.058 | 0.019 | 0.020 / 0.096 | 0.003 | 0.304 |

***Supplementary Table 3.*** *Description of the 35 metabolites with the lowest p-value after FDR correction. Name of the metabolite (Metabolite); description of their classification (Description); standardized beta (β_St_); standard error (se); 95% confidence interval (CI); p-value (p-value) and p-value after FDR correction (pFDR).*
